# Conformal growth of *Arabidopsis* leaves

**DOI:** 10.1101/048199

**Authors:** Graeme Mitchison

## Abstract

I show that Arabidopsis leaf growth can be described with good precision by a conformal map, where expansion is locally isotropic (the same in all directions) but the amount of expansion can vary with position. Data obtained by tracking leaf growth over time can be reproduced with almost 90% accuracy by such a map. The growth follows a Moebius transformation, which is a type of conformal map that would arise if there were an underlying linear gradient of growth rate. From the data one can derive the parameters that describe this linear gradient and show how it changes over time. Growth according to a conformal map has the property of maintaining the flatness of a leaf.

## Introduction

Increasingly precise information about plant growth is becoming available, and in particular the growth pattern of developing leaves and petals has been mapped in some detail [1, 2, 3, 4, 5, 6, 7, 8]. These growth patterns have been accounted for by models in which growth regulators operate within specified regions of the leaf [7], polarity fields control the predominant direction of growth [8], and gene networks regulate the succession of morphogenetic events in a combinatorial fashion [4, 5, 8].

A different type of explanation, with its roots in physics and engineering, goes back to D’Arcy Thompson and his famous book, On Growth and Form [9]. Thompson discusses leaf growth in terms of transformations extending over the whole leaf area, and points to possible underlying physical mechanisms. In a footnote ([9], p 1084), he draws attention to the resemblance of certain mappings to conformal transformations. These are transformations of the plane that preserve angles locally, and are generated by isotropic local expansion. The amount of expansion can vary with position, but at any point expansion occurs to the same extent in all directions. Conformal maps are very important in physics and engineering, because they are intimately connected with diffusion, hydrodynamics and electrical fields. There is also a close connection with complex analytic functions, which are maps from the complex plane to itself (the complex plane, or Argand diagram, represents complex numbers *x* + *iy*). Every complex analytic function is conformal and vice-versa.

A recent revival of this approach by Jones and Mahadevan [10] looks at the larger class of quasi-conformal mappings (where the expansion is not necessarily isotropic) and shows how to use them for various kinds of morphometry. Another contribution comes from Wolfram, in a footnote in his magnum opus A New Kind of Science [11], where he proposes that leaf growth might follow a conformal mapping, and that this would preserve the flatness of the leaf surface. He also observes that conformal mappings might be generated biologically by a diffusion-based mechanism, since the components of these mappings satisfy the equilibrium diffusion equation.

In a certain sense, Wolfram’s assertion about flatness is a tautology, since a conformal map is planar and a planar map will preserve flatness. The real content lies in the proposal that leaf growth is locally isotropic. If this is true, then to be planar amounts to being conformal. At first sight, it seems unlikely that leaf growth is locally isotropic; for instance, clones become markedly elongated in petals (Figure 2 in [8]) and leaves (Figure 2 in [7]).

**Figure 2:**
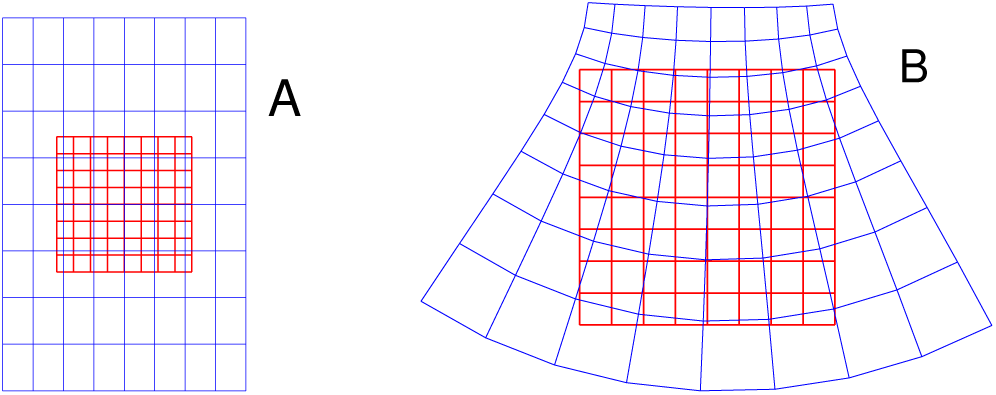
Some examples of growth maps that preserve the flatness of a leaf. A: growth is the same at all points, but can be anisotropic. B: growth is isotropic, but the growth rate can vary with position. In this case, the growth rate increases as one moves downwards, and this causes the tissue to rotate.

The surprise is that Wolfram’s assumption turns out to be largely correct. I show here that, using data kindly made available by Professor Rolland-Lagan, that the growth of *Arabidopsis* leaves approximates remarkably well to a conformal mapping, about 90% of the growth being accounted for by such a map. A fascinating question is then raised by the relationship between this kind of explanation and the gene-based interpretation discussed initially.

## Results

### The pattern of local growth

The data used here consist of a set of observations of individual *Arabidopsis* leaves, each of which has a number of beads attached to its surface that are tracked day by day. The earliest time point is day 7 after sowing, and there are altogether thirteen leaves that were tracked till day 12. As we shall see, the earliest time step, day 7 to day 8, provides the most interesting data, since uniform expansion, with a constant relative growth rate (RGR) over the entire leaf, increasingly takes over with increasing age. Figure 1 shows the displacement of beads between days 7 and 8 for several leaves.

**Figure 1:**
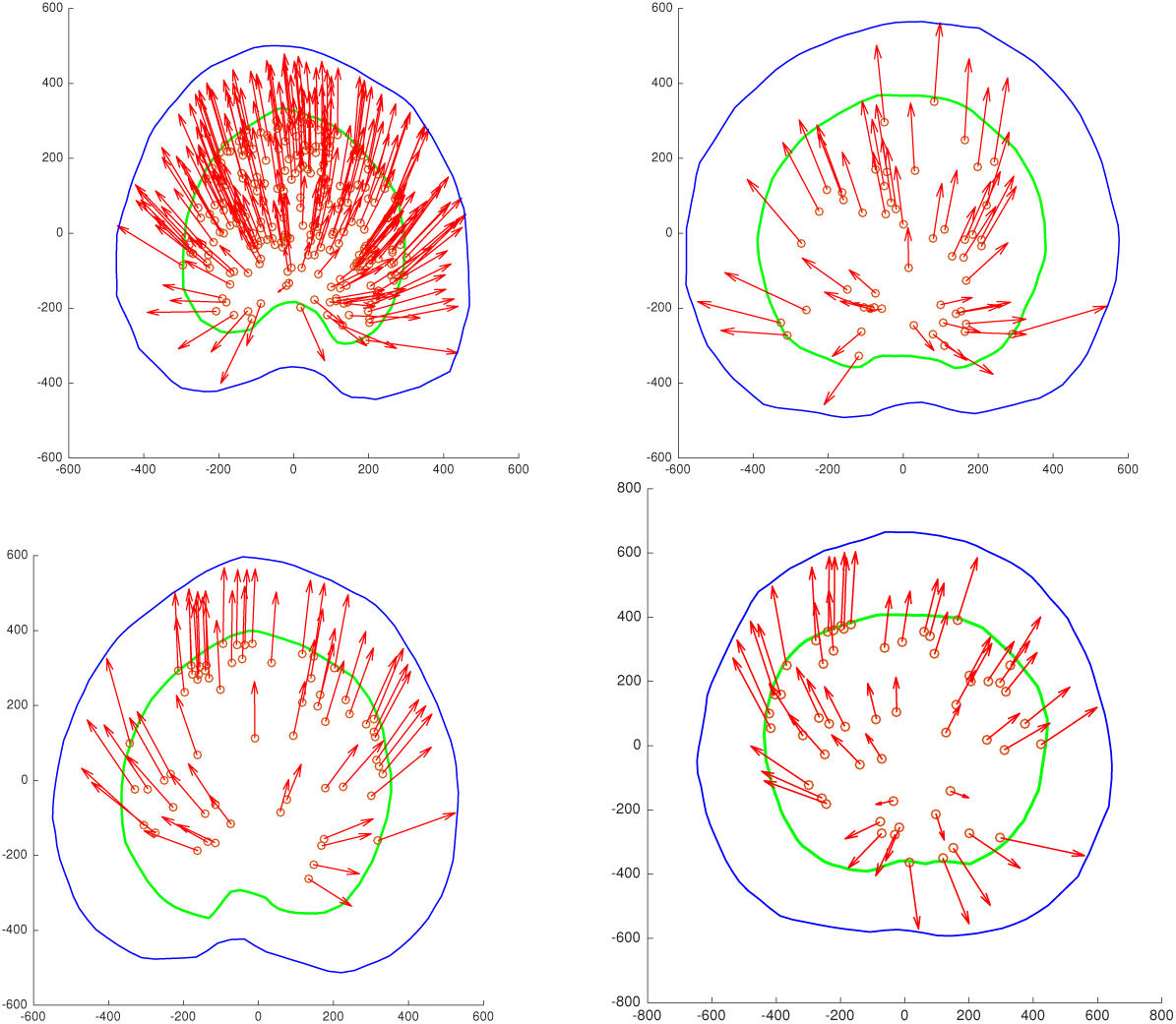
The displacement of beads between day 7 and day 8 for four leaves, the one with the greatest number of beads (leaf 1) and three others. The green outline is the margin of the leaf at day 7 and the blue outline that at day 8. The beads are shown as small circles at the start of the arrow, which ends at the position of that bead at day 8. The axes show the size of the leaves in microns. Note that the origin of the x- and, y-coordinates lies at the approximate centre of the leaves.

We would like to understand these maps at the level of individual cells and their growth rates. To do this, we need to characterise the local growth map, i.e. the way that tissues expand on a small scale. This information is captured by the derivatives of the map and in particular by the Jacobian matrix, which is the analogue, for growth between two time points, of the growth tensor [6]. If we write the map as (*x*, *y*) → (*u*(*x*, *y*), *v*(*x*, *y*)), then the Jacobian *J* is defined by

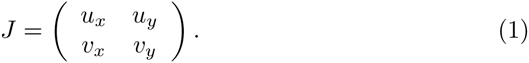

where *u*_*x*_ stands for *∂u*/*∂x*, *u*_*y*_ for *∂u*/*∂y*, and so on.

We are particularly interested in those growth rules that lead to a flat leaf surface, since many leaves are approximately flat, including the *Arabidopsis* leaves in our data set (e.g. see Figure 3 of [3]). Some of the growth rules of this type lead to a recognisable form for the Jacobian. For example, suppose that the rate of growth is the same at every point of the leaf lamina. The growth could be anisotropic (greater along one axis than the other), but the amount of growth is assumed not to vary with position. Figure 2A shows an example of such growth applied to an initial square array; each small square expands to a larger rectangle and these all have identical shape. If growth is by a factor a along the x-axis and by a factor *b* along the y-axis, the Jacobian at every point is:

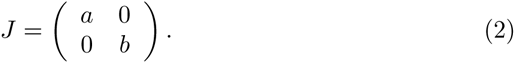

Another type of map has properties that are in a sense the opposite of the foregoing: the growth rate does not have to be constant over the leaf surface, but it is locally isotropic. Figure 2B shows an example; notice how each small square deforms into a square (approximately) because of local isotropy. Notice also that the greater growth at the bottom of the grid leads to rotation of the small squares. This is passive rotation produced by expansion of adjacent tissue [6]. The matrix for rotation through an angle *θ* is 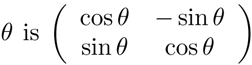. When combined with the effect of isotropic relative growth *g*(*x*, *y*), this gives rise to a Jacobian of the form

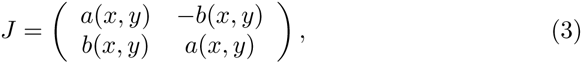

where *a*(*x*,*y*) = *g*(*x*,*y*)cos*θ*(*x*, *y*) and *b*(*x*, *y*) = *g*(*x*,*y*)sin *Θ*(*x*, *y*). Comparing Eq. 3 with Eq. 1 shows that *u*_*x*_ = *v*_*y*_ and *u*_y_ = –*v*_*x*_. These are the celebrated Cauchy-Riemann equations that define a conformal map.

**Figure 3:**
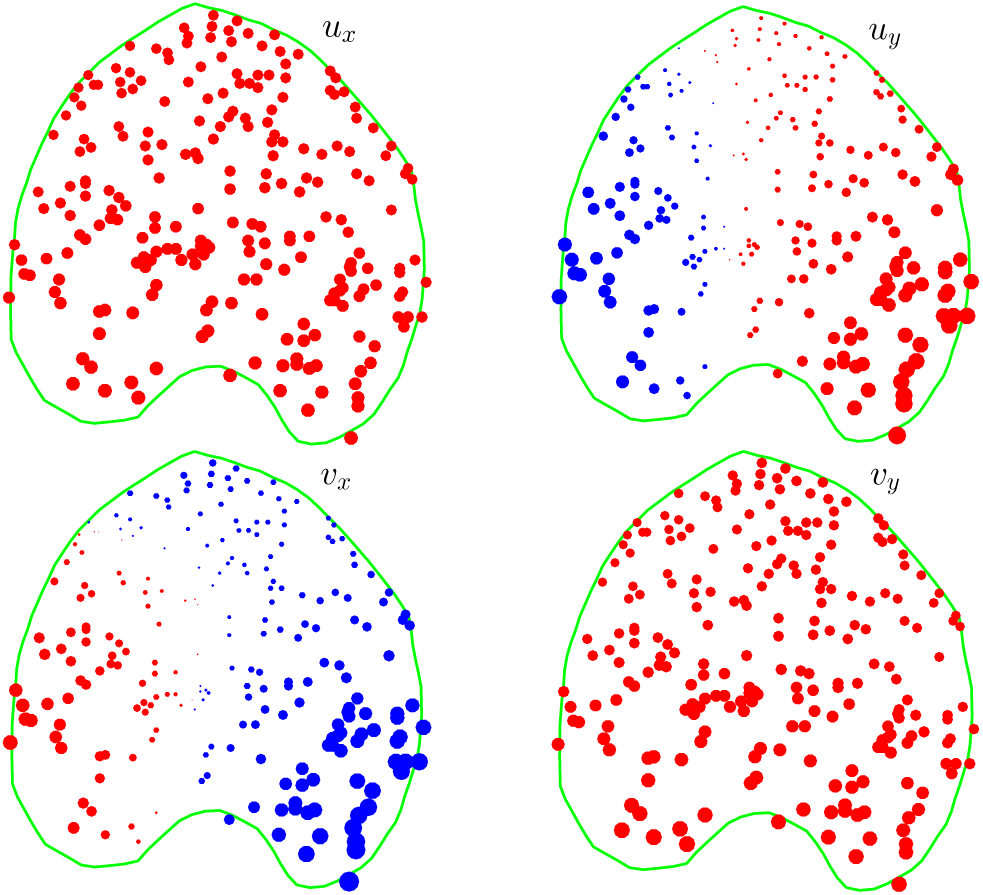
The components of the Jacobian, Eq. 3, the linear transformation that defines the local expansion and rotation due to growth. This is based on the data for leaf 1 shown in Figure 1. The radius of the beads is the (scaled) size of the derivative, with red for positive values and blue for negative. the ‘a’ components (u_x_ and v_y_) have been displayed at 1/4 the scale of the ‘b’ components (u_y_ and v_x_).

The next question is whether the Jacobian calculated for the observed leaf growth has a distinctive signature. Figure 3 shows that there is a striking agreement with the conformal pattern. The top left (*u*_*x*_) and bottom right (*v*_*y*_) derivatives are roughly equal, as expected from Equation 3, and are positive (red) because *a* = *g* cos*θ* and both *g* and cos*θ* are positive. Whereas the top right (*u*_*y*_) and bottom left (*v*_*x*_) derivatives are of opposite sign (blue is negative), also as expected from Equation 3. The fact that the signs switch around the midvein is a consequence of the symmetry of the leaf shape and the fact that growth leads to rotations in opposite directions in the two halves of the leaf. The other leaves show similar patterns, though with more noise owing to the smaller number of beads (see the last column in Table A-2).

### Fitting a conformal map to growth

In the preceding section we found that the growth map has the characteristics of a conformal map. Our strategy now will be to find the conformal map that best fits the data. Given the impressive match to the Jacobian signature, we expect this to be a rather good fit to the observed map; in other words, most of the movement of the beads will be accounted for by a conformal map. This is a strong statement about the growth of the leaves: the observed growth is mostly conformal and the basic mechanism of leaf growth must generate such a map. It also tells us something about the residual, i.e the remaining small part of the map that is not conformal. In many of the leaves this highlights regions where some kind of systematic directional growth is concentrated.

We have seen that the Jacobian signature of conformal growth is equivalent to the Cauchy-Riemann equations. These tell us that a conformal map can be regarded as a complex analytic function (see Eqs A-3 and A-4). Given this, our strategy will be to interpret the plane in which a leaf grows as the complex plane. This means that a point with coordinates (*x*, *y*) is regarded as the complex number *z* = *x* + *iy*. We can then look at any function of *z*, e.g. a polynomial *a* + *bz* + *cz*^2^, as a map from the complex plane to itself, and we can ask how well this approximates the growth of the leaf - regarding this also as a map of the complex plane to itself. By varying the coefficients *a*, *b* and *c* (which are themselves complex numbers) we can find the polynomial that minimises the error in predicting the positions of the beads.

Consider the linear function *f*(*z*) = *a*+*bz*, where *z* = *x*+*iy*, and *a* = *a*_0_ + *ia*_1_ and *b* = *b*_0_ + *ib*_1_ are complex constants. This map takes the point (*x*, *y*) to (*u*, *v*), where *u* = *a*_0_ + *b*_0_*x* – *b*_1_*y* and *v* = *a*_1_ + *b*_0_*y*+*b*_1_*x*. This amounts to a shift of origin to the point *a* in the complex plane, a rotation by arg*b* = arctan(*b*_1_/*b*_0_), and a constant RGR everywhere on the leaf of 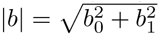. Thus a linear complex map can capture the relative positioning and orientation of the leaf between successive days, which have to do with the experimental set-up; it can also account for a constant RGR, which is a parameter with biological significance.

Thus the best-fitting linear complex map, found by least squares fitting of the bead data points, gives us an estimate of both the experimental parameters and the intrinsic uniform isotropic growth. The difference between the final bead position predicted by this linear model and the true position, i.e. the linear model *residual*, indicates how much of the growth involves variation of growth rate with position in the leaf, with consequent rotation of the tissue, as depicted in Figure 2B. The size of this residual can be measured by summing the lengths of the residual vectors and dividing this by the sum of the total bead displacement observed experimentally. This *normalised residual* has an average value over all the leaves of 0.33; see Table A-2. Equivalently, one can say that 67% of the displacement is accounted for by a linear model.

The distribution of this residual over the leaf lamina is shown in Figure 4. This is very striking: as one follows a path round the perimeter of the leaf, the residual vectors rotate *twice as fast* as the radial vector. This is what is expected of a quadratic complex function, as shown in Figure 5. Note that the origin of the leaf’s coordinate system is in the approximate centre of the leaf, and that of the complex quadratic lies at the centre of the circle.

**Figure 4:**
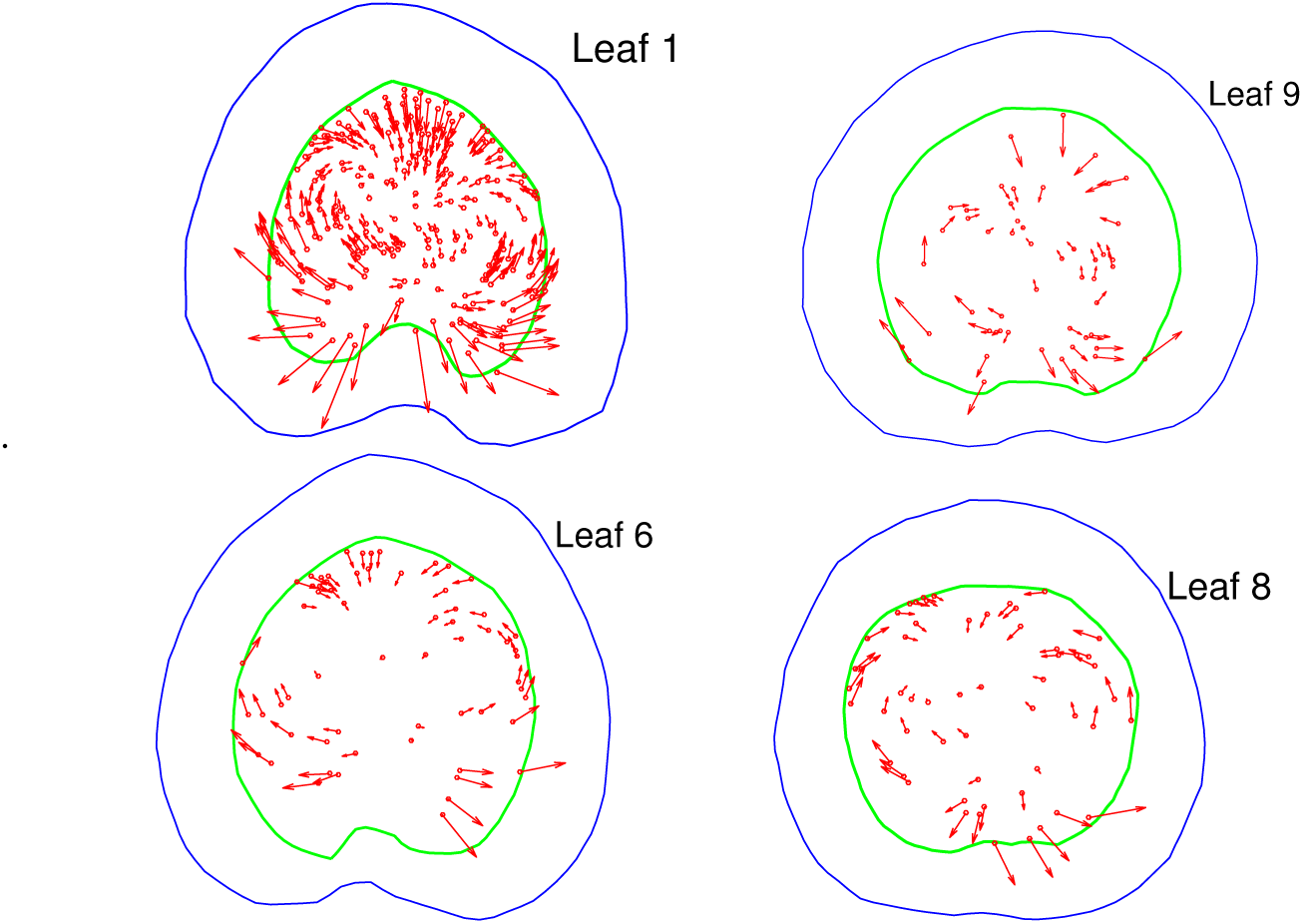
The residual movement of beads after subtracting the best-fitting linear map, for four leaves and the time period 7-8 days. Note the rotation of the arrows at twice the speed of the radius, and compare with the quadratic function in Figure 5

**Figure 5:**
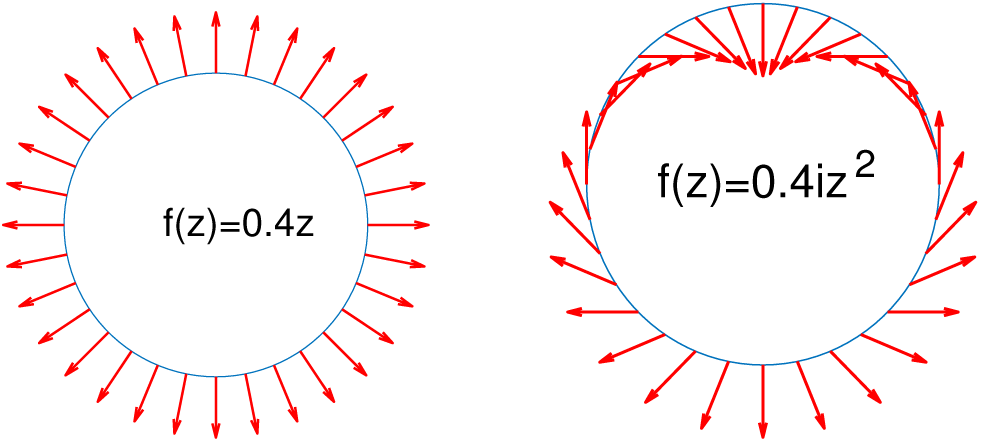
Two complex functions, represented by arrows: the linear function, f (z) = 0.4z and the quadratic one f (z) = 0.4iz^2^.

Given this pattern in the residual, we expect to get a better fit to the data with a quadratic complex polynomial *f* (*z*) = *a* + *bz* + *cz*^2^, and this is indeed the case, as shown by the substantially smaller normalised quadratic residuals in Table A-2, with an average of 0.14, (86% of the displacement accounted for). Adding a cubic term, *f*(*z*) = *a* + *bz* + *cz*^2^ + *dz*^3^, gives a further small improvement, with 0.11 residual or 89% of the displacement accounted for; see Table A-2. Additional higher power terms give negligible improvement.

In addition to fitting a polynomial, one can also fit a Mobius transformation, which has the form

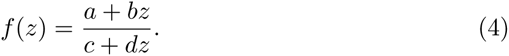

Möbius transformations have the attractive property that the composition of two of them - i.e. following one by the other - is again a Mobius transformation. Furthermore, they have a matrix representation that is very useful for interpolating between the observations, as will be discussed later. It turns out that one gets almost as good a fit from a Möbius transformation as from a cubic, with an average residual of 0.12, or 88% of displacement accounted for; see Table A-2.

To summarise so far, a linear conformal map, which is equivalent to constant RGR, can account for about 67% of the displacement of beads, whereas a nonlinear conformal map, cubic or Mobius, accounts for 88-89% of the displacement (these are figures averaged over all leaves, for days 7 to 8). There is therefore a substantial nonlinear component to the conformal growth.

The residual of the best-fitting conformal map highlights regions of anisotropic or directional growth. As can be seen from Figure 6, there is considerable variation between individual leaves, some having marked anisotropic basal growth and others having very little. Basally localised anisotropic growth has previously been observed in averaged leaf data, e.g. Figure 7 in [2]. What is new here is the large amount of individual variability in this anistropic component.

**Figure 6:**
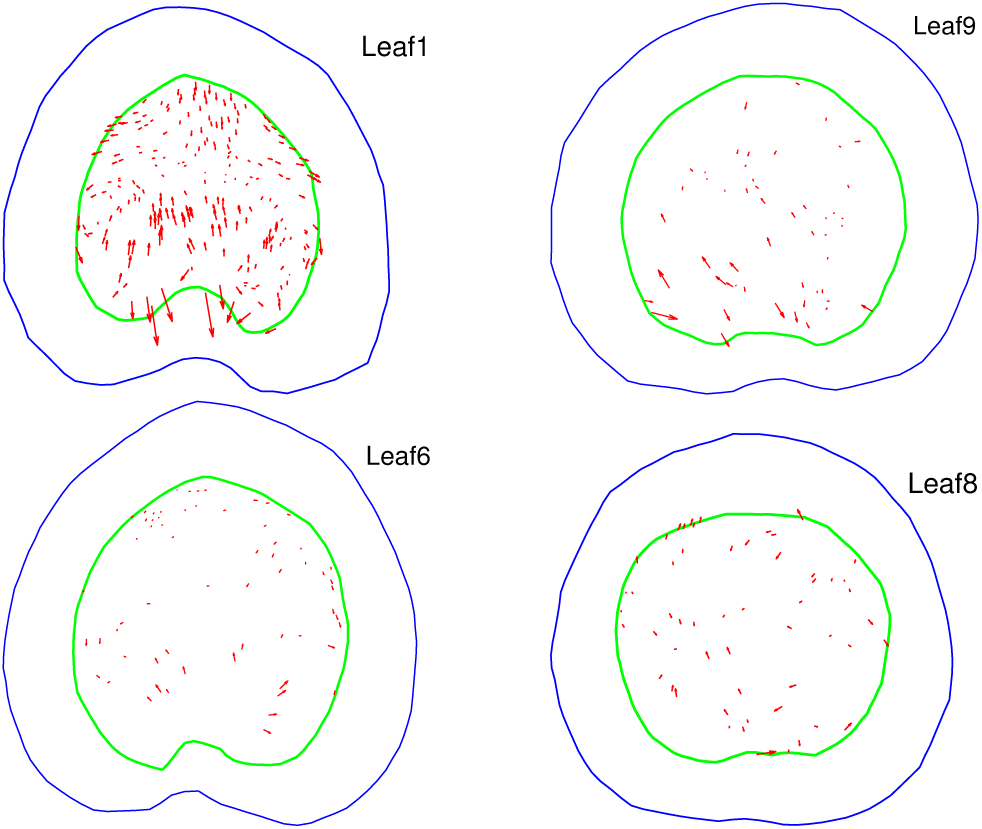
The residual from the best fitting complex polynomial, for four leaves from days 7-8. The small circles representing beads have been removed so that the arrows can be seen more clearly.

The leaves also show a strong basal bias in the *isotropic* growth rate, which can seen in the best-fitting conformal map. Unlike anisotropic growth, this pattern is found in all the leaves. This is well documented in the literature [7, 3], though fitting a conformal map clarifies the distinction between oriented and isotropic components in this gradient.

The data set allows one to follow the pattern of residuals for five successive time steps, beginning with 7-8 days. Figure 7 shows residuals averaged over all leaves: the linear residual gets steadily smaller, implying that the component of constant RGR increases steadily, so that some 85% of the growth is constant expansion by days 11-12. The pattern of linear residuals for an individual leaf (leaf 1) over five successive days is shown in Figure 8, and it tells the same story, with the initially large residuals at the apex and base of the leaf gradually diminishing.

**Figure 7:**
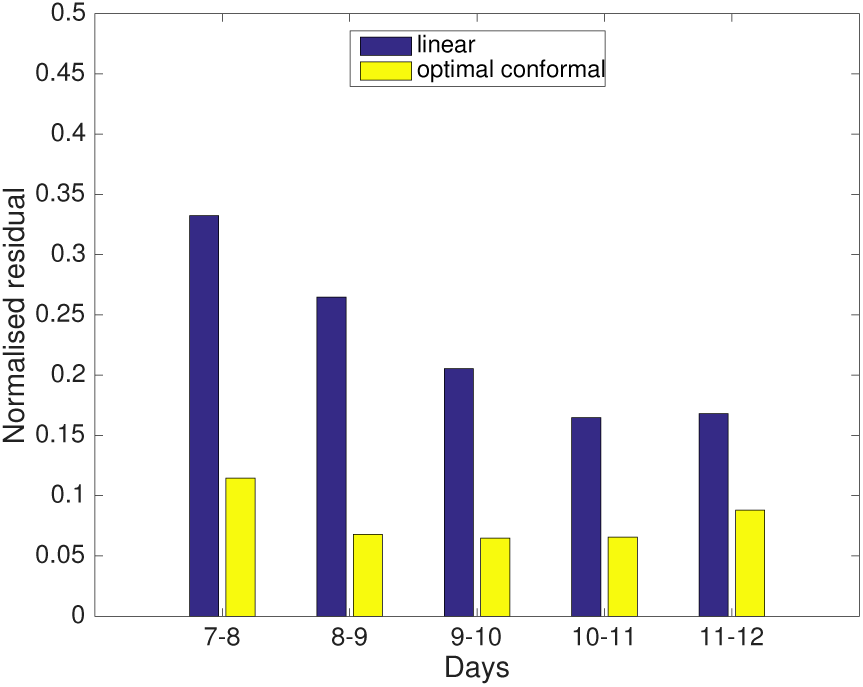
Change in the average residuals (linear and cubic) over time, showing the decrease in the linear residuals, and hence the increasingly large component of constant RGR, as the leaves age.

**Figure 8:**
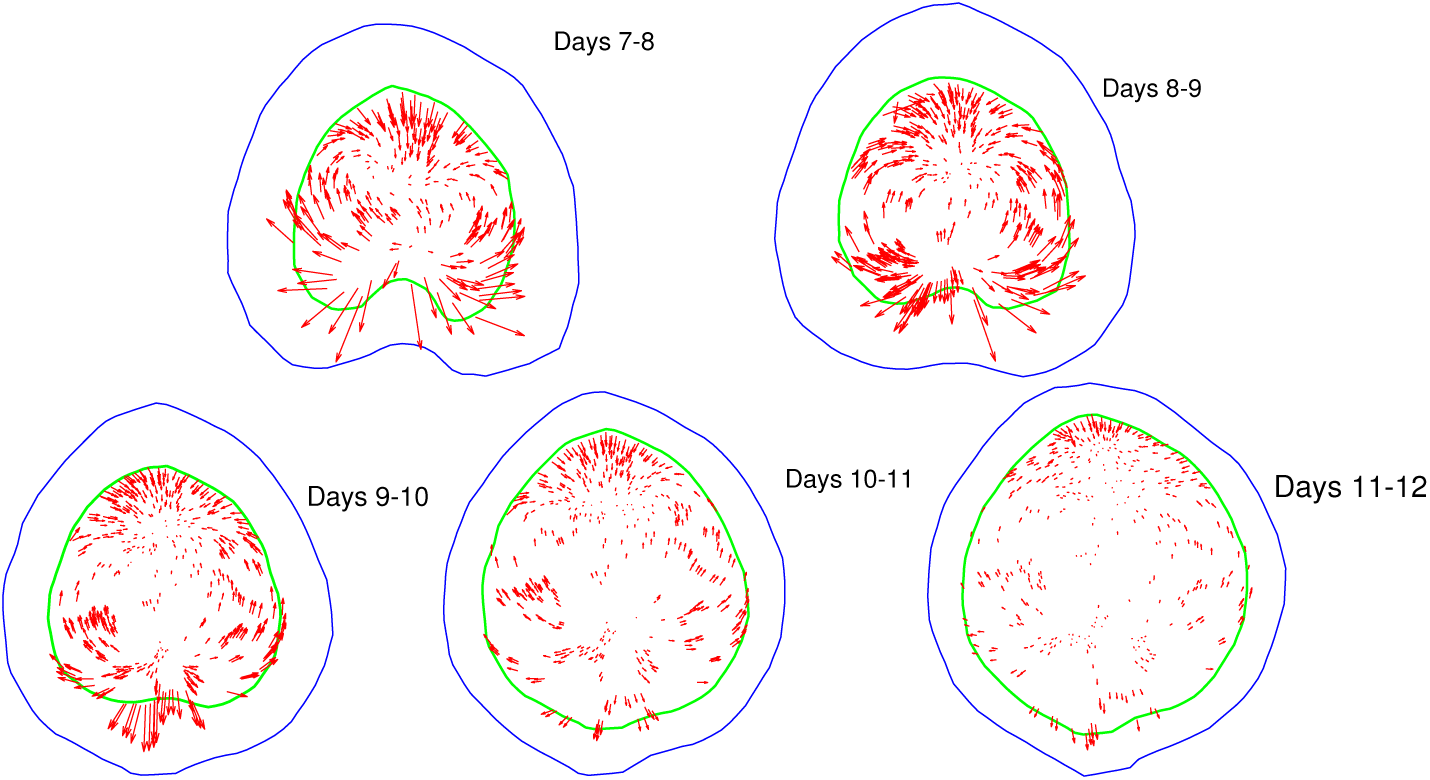
The linear residual for leaf 1 from day 7 to day 12.

### The biological meaning of conformal growth

A conformal map is a very special type of map, and it is natural to ask how it could be generated biologically. Wolfram’s answer [11] is that conformal growth will occur precisely when the RGR has the special property of being a harmonic function, i.e. when it satisfies the equilibrium equation for diffusion. He also proposed that the RGR might therefore be specified by the concentration of a diffusing signal molecule, but I will point out some difficulties with this proposal in due course.

Now it turns out that for the leaves in our data set the conformal component of their growth is well approximated by a Möbius function, Eq. 4, and that the underlying RGR that generates these functions is of a particularly simple kind, namely a linear gradient. All that is required, therefore, is that cells set their RGR - their rate of isotropic expansive growth - according to a linear gradient at any moment in time. A linear gradient is a harmonic function, but a very special one.

Let us run through the steps that lead to this conclusion. A Mobius function has the attractive property that one can ‘wind back the clock’ and recreate intermediate steps in the growth pattern. This is possible because composing two Mobius transformations is equivalent to multiplying their matrices. Thus given an observed map over some time period (24 hours for the data used here), one can take a fractional power of the associated matrix and estimate the map for a shorter time period than the original 24 hours. Figure 9 illustrates the method by taking the square root of the matrix *A* for the best-fitting Mobius transformation for leaf 4 between days 7 and 8; the matrix 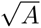 gives an estimate of growth in half the period of observation, i.e. 12 hours.

**Figure 9:**
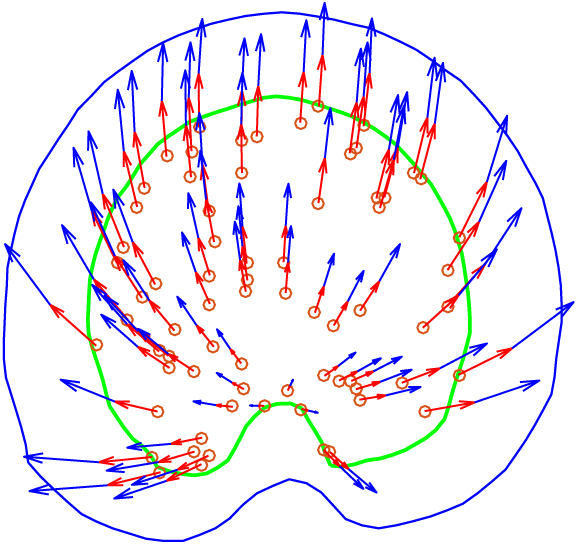
Factorising the growth into two 12 hour periods, using the Mobius transformation with matrix 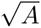, where A represents the best fit for the 24 hours between days 7 and 8 (leaf 4)· The composition of the two maps, i.e. the result of following the red arrows and then the blue arrows, is equivalent to the 24 hour best fit.

By taking smaller fractional powers one retrieves the growth pattern for shorter times, and in the limit this enables one to calculate the infinitesimal generator (Eqns A-10, A-11 and A-12), and hence the distribution of the RGR everywhere on the leaf. A straightforward calculation, Eq. A-14, shows that the RGR distribution is linear, and one can estimate the constants that determine this linear gradient from the bead movements. Table 1 gives these constants for leaf 1 over five successive day periods. There is a steady decrease in the average growth rate with time, (*b*_0_ – *c*_0_), and also a decrease in the slope in the direction of the leaf axis, *d*_1_, which accords with the decreasing nonlinear terms shown in Figures 7 and 8. Figure 10A and B show what the gradients look like for this particular leaf.

**Table 1:** The constants specifying the linear gradient in the RGR for leaf 1 over five successive day-lengths. The gradient is given by Eq. A-14 as RGR(x,y) = (b_0_ – c_0_) – 2d_0_x – 2d_1_y. Growth is over 24 hours, so the units for b_0_ – c_0_ are per day, and for d_0_ and d_1_ are per millimetre per day.

| days | $b_0 - c_0$ | $d_0$ | $d_1$ |
| --- | --- | --- | --- |
| 7-8 | 0.4457 | 0.038 | 0.574 |
| 8-9 | 0.4362 | 0.015 | 0.271 |
| 9-10 | 0.3391 | 0.004 | 0.120 |
| 10-11 | 0.2686 | -0.006 | 0.051 |
| 11-12 | 0.2099 | 0.003 | 0.026 |

**Figure 10:**
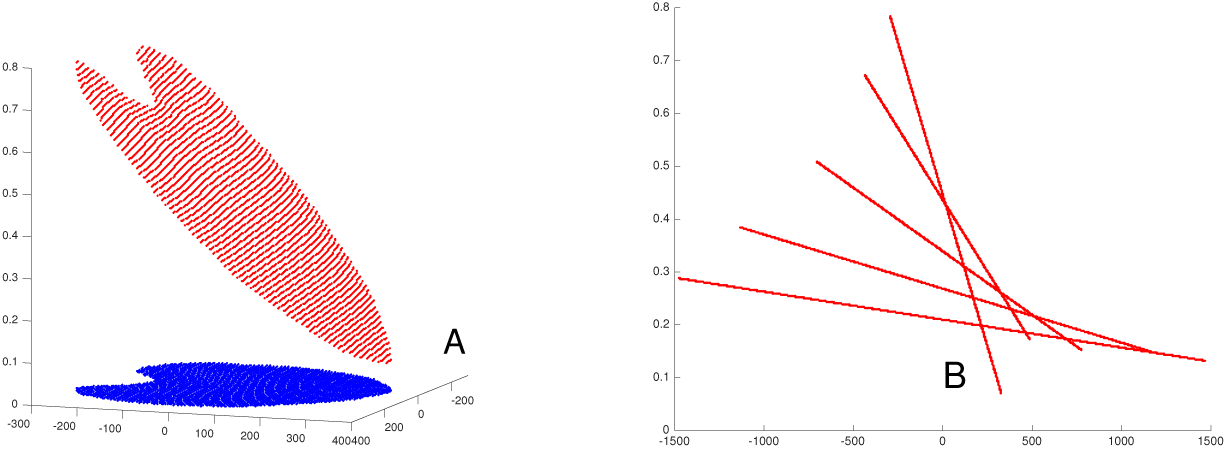
A: The linear gradient in RGR. Its minimum value is at the distal tip of the leaf, where growth is slowest, and its largest values at the base. B: The gradual flattening of the gradient in RGR with successive days. Each line represents the gradient along the midvein.

## Discussion

The growth patterns of the leaves studied here have a significant nonlinear component (Figure 4), yet it turns out that they can be explained by a linear gradient of growth rate (Figure 10 A and B). It is interesting to ask how this linear gradient could be generated. Linear gradients of a diffusible signal molecule were originally postulated as the underlying mechanism for pattern formation [12, 13], and this seems natural for certain geometries, e.g. a segment in an insect embryo, since diffusion will interpolate in a linear fashion between a source and sink. However, a linear diffusion gradient in a leaf can only arise in a somewhat contrived way, since the outline of the leaf is irregular, and there needs to be a special layout of sources and sinks to ensure linearity (see Figure 11). Furthermore, the gradients that have been found are not linear; in the case of Bicoid, the gradient is exponential [14, 15, 16].

**Figure 11:**
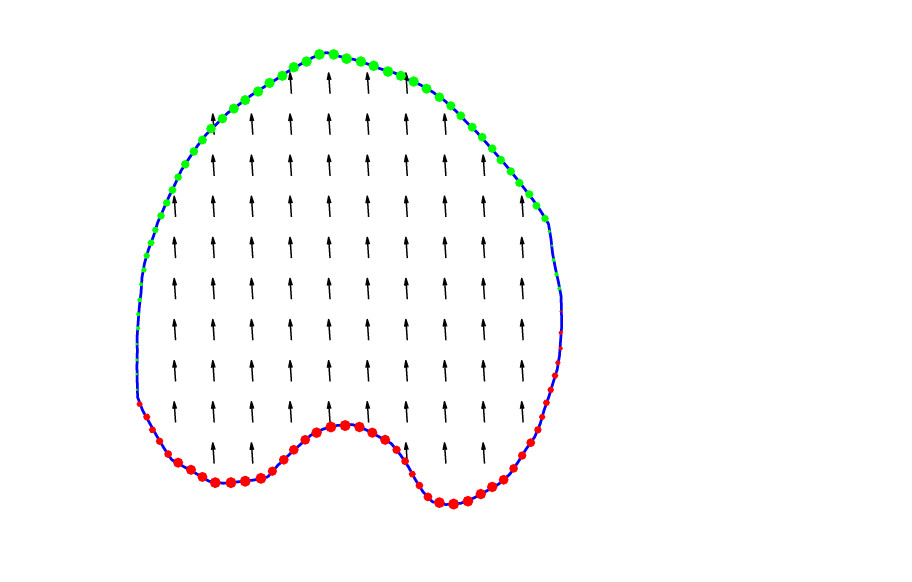
The flux pattern associated to a linear gradient set up by diffusion, and, the distribution of sources (red) and sinks (green), calculated for the outline of leaf 1, that would be needed to generate the gradient.

Thus Wolfram’s [11] proposal of read-out of the RGR from an equilibrium diffusion gradient seems implausible here, though one abandons this idea with regret since it would have meant that any source or sink arrangement on the margin of a leaf would create a signal distribution that was guaranteed to produce a conformal map and hence a flat leaf. If this is not the explanation, then it is an intriguing question what mechanism leads to a linear gradient of RGR over the leaf lamina. It also invites investigation of other leaves to see if their growth is also conformal and whether it is also well approximated by a Mobius transformation.

There is already a fairly detailed analysis of leaf growth in terms of developmental programs [1, 2, 3, 4, 5, 6, 7, 8], and it would be interesting to bring this into register with the fact that growth is largely conformal. Can the action of the hypothetical factors PGRAD and LATE in the leaf shape model of Kuchen et al. [7] be explained by the linear growth rate in the conformal picture and its diminishing slope with time (Table 1 and Figure 10)? One difference is that PGRAD levels are assumed to be inherited by lineage and therefore to deform with the growth of the tissue [7], whereas the linear RGR in our conformal picture changes dynamically with time so as to maintain its linearity. There is also no indication of a distinction between lamina and midvein growth (formalised by the factors LAM and MID [7]) in the residuals from the best-fitting conformal map (e.g. Figure 6). However, this may just be a matter of resolution, since the domain of operation of MID is quite narrow.

It would also be interesting to understand how the mechanism proposed to explain indentations in *Arabidopsis* leaves [17, 18] fits with a conformal viewpoint. Is this a developmental subroutine added onto the conformal growth plan of the lamina, or is it integrated into the underlying RGR distribution to make a complete conformal map?

## Acknowledgements

I thank Anne-Gaelle Rolland-Lagan for allowing me to use her data, Benoit Tremblay for help with the Matlab package that allows one to manipulate the data, Konrad Wagstyl for an introduction to Matlab programming, and Julie Ahringer, Dennis Bray, Jeremy Gunawardena and Ottoline Leyser for constructive comments on the manuscript.

## Appendix

### Data and analysis programs

The two papers from Prof. Rolland-Lagan’s laboratory, [2, 3], give a thorough description of growth patterns and point the reader towards a website (http://hdl.handle.net/10393/30401) where the bead data for individual leaves are available, and also a package of Matlab programs for leaf shape analysis. Leaves are referred to by pot and plant numbers in their data sets, and Table A-1 shows how this relates to the numbering used here.

**Table A-1:**
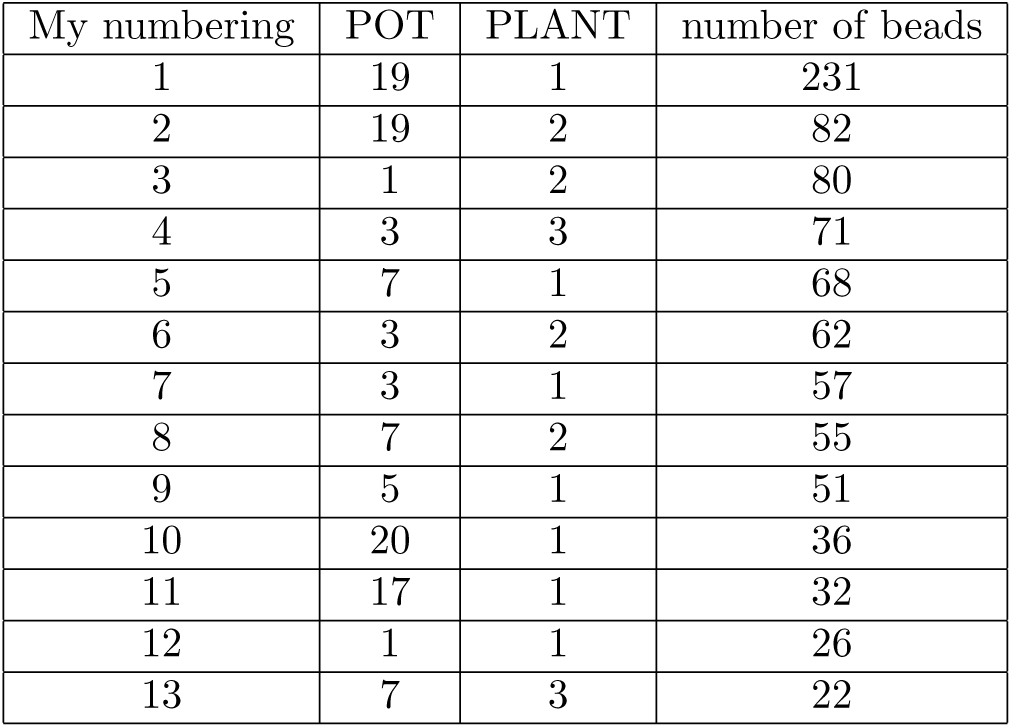
The numbering of leaves used here (in order of decreasing number of beads), and the numbering in [2, 3], defined, by the pot and plant. Also given is the number of beads on each leaf.

### Calculating and interpreting the Jacobian

To compute the Jacobian (Eq. 1) at each bead we need to estimate the derivatives *u*_*x*_, *u*_*y*_, *v*_*x*_, and *v*_*y*_. For this we need a map (*x*,*y*) → (*u*(*x*, *y*), *v*(*x*, *y*)) that interpolates smoothly between the values at beads. Following [1], we assume that *u* and *v* are each third-order polynomials in *x* and *y*, and choose values of the polynomial coefficients that minimise the sum-of-squares error, *S*. If the coordinates of bead *k* at time 1 are 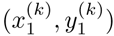 and at time 2 are 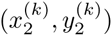, the error is given by:

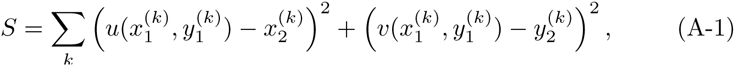

where the index *k* runs over all beads. The minimisation is a linear problem that can be done very rapidly by linear algebra functions in Matlab.

No assumption is made about the form of the polynomials, and there is no presumption that they lead to a conformal map. However, as Figure 3 shows, for leaf 1 the map shows the diagnostic sign pattern of a conformal map. To quantify this for all the leaves, we would like to test how exactly the Cauchy-Riemann equations *u*_*x*_ = *v*_*y*_, *u*_*y*_ = – *v*_*x*_ hold. The normalised inner product *R* can be used as a measure. If we take

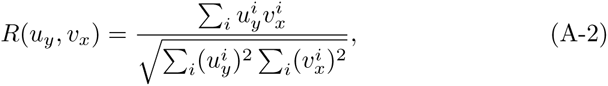

where *i* runs over all beads, then *R*(*u*_*y*_, *v*_*x*_) = 1 means that *u*_*y*_ = *v*_*x*_ for all beads and *R*(*u*_*y*_,*v*_*x*_) = –1 means that *u*_*y*_ = –*v*_*x*_ for all the beads. The last column of Table A-2 shows that *R*(*u*_*y*_,*v*_*x*_) is close to -1 for all but leaves 5, 11 and 12, and leaves 5 and 11 have the largest residuals from the best-fitting conformal map. We can also take the equivalent measure *R*(*u*_*x*_,*v*_*y*_) for *u*_*x*_ and *v*_*y*_, but in this case the measure is very close to +1 (larger than 0.98) for all leaves, so we conclude that *u*_*x*_ = *v*_*y*_ holds more closely for most beads.

#### Fitting conformal maps

Conformal maps are fitted by least squares. For instance, a quadratic map *f*(*z*) = *a* + *bz* + *cz*^2^, with *z* = *x* + *iy*, can be written in the form *f*(*z*) = *u*(*x*, *y*) + *iv*(*x*, *y*), where

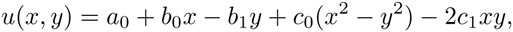

and

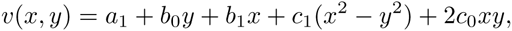

with *a* = *a*_0_ + *ia*_1_, *b* = *b*_0_ + *ib*_1_, *c* = *c*_0_ + *ic*_1_. then we find the values of *a*_0_, *a*_1_, *b*_0_, *b*_1_, *c*_0_, *c*_1_ that minimise the sum of squares error given by Eq. A-1.

The Möbius transformation is fitted by writing the function *f*(*z*) = (*a* + *bz*)/(*c* + *dz*) as a polynomial series, viz. *f*(*z*) = (*a*/*c* + *b*/*c*)(1 – *dz*/*c* + *d*^2^*z*^2^/*c*^2^ – …), and comparing its terms up to quadratic order with those of the best-fitting cubic. This determines *a*, *b*, *c*, *d*, up to an irrelevant shared factor. The cubic term in the expansion is then found to be quite closely approximated by that of the best-fitting cubic (which is in any case a very small term). In other words, the best-fitting cubic is already quite close to a Möbius transformation. This can be seen in the small changes in the residuals in Table A-2.

**Table A-2:** Normalised residuals after least-squares fitting to linear, quadratic or cubic complex polynomials. The normalised residual is defined as Σ_i_|v_i_ – l_i_|/Σ_i_|v_i_|, where v_i_ is the vector between bead positions on successive days, 1_i_ is the vector predicted by the complex polynomial, and i runs over all beads. Also shown in the measure R(u_y_,v_x_) of accuracy with which the Cauchy-Riemann equation u_y_ = ‒v_x_ holds. If R = –1, this equation holds exactly for every bead. As can be seen it is negative (except for one leaf) and close to -1 for some leaves.

| Leaf | linear residual | quadratic residual | cubic residual | Möbius residual | $R(u_y, v_x)$ |
| --- | --- | --- | --- | --- | --- |
| 1 | 0.289 | 0.124 | 0.101 | 0.103 | -0.99 |
| 2 | 0.449 | 0.202 | 0.141 | 0.148 | -0.89 |
| 3 | 0.381 | 0.158 | 0.135 | 0.130 | -0.75 |
| 4 | 0.376 | 0.164 | 0.138 | 0.141 | -0.74 |
| 5 | 0.406 | 0.209 | 0.165 | 0.178 | -0.40 |
| 6 | 0.247 | 0.083 | 0.052 | 0.065 | -0.97 |
| 7 | 0.325 | 0.117 | 0.110 | 0.112 | -0.88 |
| 8 | 0.348 | 0.145 | 0.092 | 0.116 | -0.87 |
| 9 | 0.301 | 0.121 | 0.099 | 0.107 | -0.79 |
| 10 | 0.261 | 0.123 | 0.097 | 0.118 | -0.81 |
| 11 | 0.360 | 0.178 | 0.180 | 0.193 | 0.18 |
| 12 | 0.253 | 0.101 | 0.101 | 0.105 | -0.07 |
| 13 | 0.327 | 0.104 | 0.080 | 0.088 | -0.90 |
| average | 0.333 | 0.141 | 0.115 | 0.123 |  |

#### Complex functions and their derivatives

Let *f* be a complex analytic function. This means that *f* is differentiable as a complex function (as are all the functions one is likely to encounter in the present context). We can write the function in terms of its real and imaginary parts as *f*(*x*, *y*) = *u*(*x*, *y*) + *iv*(*x*,*y*), where *x* + *iy* is the point with coordinates *x*, *y* in the complex plane. Calculating the derivative of *f* by a real *δx* gives *f′* = *u*_*x*_ + *iv*_*x*_ (where *u*_*x*_ denotes *∂u*/*∂x*, etc.), whereas making the calculating with an imaginary *iδy* gives *f′* = –*iu*_*y*_ + *v*_*y*_. Equating these two definitions yields the Cauchy-Riemann equations,

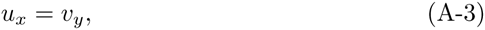

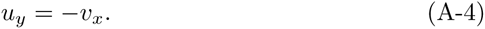

These in turn imply that

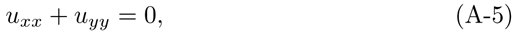

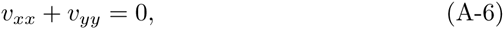

(where *u*_*u*_ = *∂*^2^*u*/*∂x*^2^, etc.), implying that both *u* and *v* satisfy Laplace’s equation, which is the equation for the equilibrium distribution of a diffusing substance [19]. A function satisfying this equation is called a *harmonic* function. Thus both the real and imaginary parts of *f* are harmonic. Conversely, given a harmonic function *u*, the Cauchy-Riemann equations can always be solved to give *v*, and this yields a conformal map with real and imaginary parts *u* and *v*.

#### The relative growth rate of a conformal map family

We begin with an assumption, that is only an approximation to biological reality, that growth results from the repeated iteration of the same map that operates over a short time period. Thus we define the map *f*_1/*n*_ to be the map whose n-fold composition is the map *f*(*z*) observed over some time interval (e.g. the 24 hours of the observations here). In other words

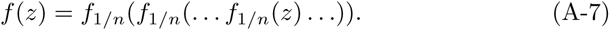

We assume that such a map *f*_1/*n*_ exists for arbitrarily large n (i.e. arbitrarily short times). The reason this is not biologically fully plausible is that the map is likely to change as the leaf grows: what we are calling *f*_1/*n*_ will not be quite the same map at the beginning and end of a sequence of iterations. However, the simplifying assumption makes the problem mathematically tractable, and is enshrined in the concept of a semigroup of maps, where *f*_*s+t*_(*z*) = *f*_*s*_(*f*_*t*_(*z*)), for all positive *s*,*t* [20].

For large *n*, the map *f*_1/*n*(*z*)_ is close to the identity, and can therefore be written approximately as

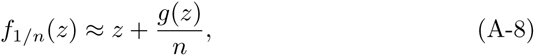

where *g*(*z*) is called the *infinitesimal generator* of the semigroup [20]. If *f*(*z*) is conformal, so is *f*_1/*n*_(*z*) and so is the infinitesimal generator. Now define the relative growth *RG*_*f*_(*z*) for the map *f*(*z*) at the point z as |*f*′(*z*)|. The relative growth *rate*, *RGR*(*z*), which is what we want to find, is the temporal derivative of *RG*(*z*). If *g*(*z*) has real and imaginary parts *h* and *k*, respectively

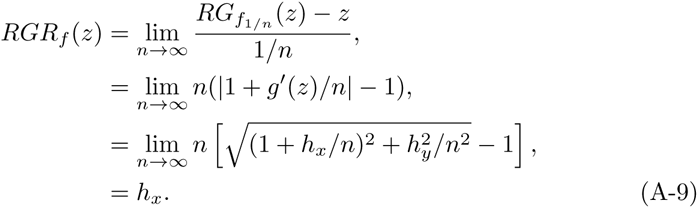

Now *h*_*x*_, which is the real part of the analytic function *g*′(*z*), is harmonic. So the growth rate could be specified by the concentration of a substance at diffusive equilibrium. These steps can be reversed, since any harmonic function can be written as the real part of some complex function *k*(*z*) and we can take the conformal infinitesimal generator to be *g*(*z*) = ∫ *k*(*z*). Since composition of conformal maps is conformal, the resulting growth is conformal. So specifying the growth rate by the concentration of a diffusible substance produces a conformal map, a conclusion reached by a different argument, far more tersely and elliptically, in [11].

#### The relative growth rate of a Mobius transformation family

Calculating the infinitesimal generator is easy for a Mobius transformation *f*. If *A* is the matrix representing *f*, then *A*^*1/n*^ is the matrix representing *f*_1/*n*_ (see Figure 9 for an illustration of *f*_1_/_2_). In the limit of large *n*,

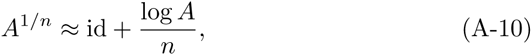

where log *A* is the matrix logarithm of *A*. Writing

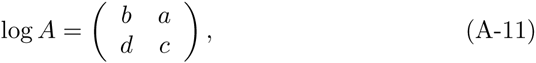

and converting the matrix *A*^1/*n*^ back into a Mobius transformation gives (neglecting terms in (1/*n*)^2^ or higher powers)

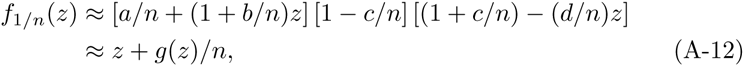

where

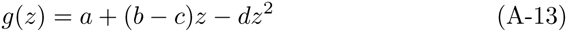

Thus the infinitesimal generator *g*(*z*) is a quadratic polynomial. Eq. A-9 now gives

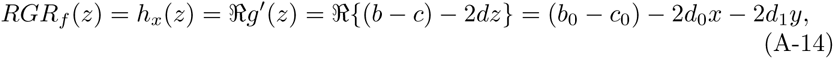

where *b*, *c*, *d* are the entries in log(*A*) defined above with *b* = *b*_0_ + *ib*_1_, *c* = *c*_0_ + *ic*_1_, *d* = *d*_0_ + *id*_*1*_. Thus three real parameters are needed to define the growth.

